# General formulation of Luria-Delbrück distribution of the number of mutants

**DOI:** 10.1101/019869

**Authors:** Bahram Houchmandzadeh

## Abstract

The Luria-Delbrück experiment is a cornerstone of evolutionary theory, demonstrating the randomness of mutations before selection. The distribution of the number of mutants in this experiment has been the subject of intense investigation during the last 70 years. Despite this considerable effort, most of the results have been obtained under the assumption of constant growth rate, which is far from the experimental condition. We derive here the properties of this distribution for arbitrary growth function, for both the deterministic and stochastic growth of the mutants. The derivation we propose is surprisingly simple and versatile, allowing many generalizations to be taken easily into account.

## I. INTRODUCTION

The Darwin-Wallace theory of evolution rests upon mutation of living organisms and their selection. In their seminal article [1], Luria and Delbriück (LD) described an experiment demonstrating the randomness of mutational events *before* the selection process. The experiment consists of growing *C* cultures of bacteria in parallel in identical environments, beginning with a small number *N*_0_ (typically 10^3^) in each batch. After a sufficient growth period, the cultures saturate and the number of bacteria reaches *N* (typically 10^9^ − 10^10^). Each culture is then tested against an antibacterial agent, a phage virus in the LD case, and the number of surviving bacteria arising from mutation in the cultures is counted by a plating method. If *C* is large, the probability *P* (*m*) of having *m* mutants can be experimentally determined; in practice, *C* cannot be large and therefore only statistical quantities such as the mean and the variance of the number of mutants can be estimated.

The LD experiment has spurred a large interest and many authors have developed increasingly refined models to estimate statistical properties of the random variable *X* of the number of mutants, such as its cumulant/probability/moment generating functions (cgf, pgf, mgf), from which the mutation rate or probability can be estimated. The pioneering authors were Lea and Coulson[2], Armitage [3], Bartlett[4], Crump and Hoel[5], Mandelbrot[6], Sarkar, Ma and Sandri[7] who set the LD distribution on a solid mathematical ground, generalized the model to take into account stochastic growth of the mutant and the wild type, and developed algorithms to estimate the mutation rate. The works of these and other authors have been reviewed in an elegant article by Zheng[8] which also contains original results and corrections of some of the errors contained in the previous works. These investigations have been extended during the last 15 years by authors such as Angerer[9], Dewanji et al[10] and Ycart[11]. A description of some of these more recent works will be given in the following sections.

The fundamental LD experiment is now currently used to estimate mutation rates in various setups such as antibiotic resistance or experimental investigation of the evolutionary process[12–14].

However, nearly all of the existing computations have focused on the exponential growth (either deterministic or stochastic) of the WT and mutant bacteria, although Dewanji et al.[10] have extended these results to Gompertz growth. The reason behind this choice is that in these formulations, an explicit expression for the number of wild type (WT) cells as a function of time is needed in order to compute the statistical properties of the number of mutants.

The assumption of constant growth rate is however too restrictive. Experimentally, the growth is never exponential but follows a Monod curve[15] the growth rate is not constant, but begins with a value close to zero (called the lag phase), increases gradually to a maximum value and then decreases as the number of bacteria increases, to finally reach zero when the culture is saturated. Various functions (logistic, Gompertz, Richards, Stannard, …) are used in the literature to model the growth curve and their relevance has been studied in depth by Zwieteting et al. [16].

The *real time* however in not the relevant independent variable in terms of which the system may be described. The WT population grows from an initial number of cells *N*_0_ to reach a final value *N*. Each time a WT cell divides, there is a small probability *ν* that a mutant having the desired trait (phage or antibiotic resistance) appears. It does not matter how much time the system spends between WT population size *n* and *n* + 1, but only the fact that once a division has taken place, a mutation may appear. For the mutants growing in the same environment as the WT, their growth curve will be similar (but not necessarily equal) to that of the WT. The only quantities that are indeed measured in an experiment are *N*_0_, *N* and the number of mutants. Even though the growth curve can theoretically be measured, its determination is cumbersome and, as we will see, not relevant.

In this paper, we shall use the WT population size *n* as the independent variable. It appears that this formulation of the LD distribution is surprisingly simple and applies to any growth curve for the bacteria, including of course exponential growth. The formulation is versatile and can take into account many generalizations, some of which are considered in this article.

This article is organized as follows: in the next section, we present the basic concept for the simple case of deterministic growth of both WT and mutant bacteria, where only mutations apparitions are random. We present some generalizations, such as different growth rates for WT and mutants and non-constant mutation probability. In the following section, we generalize the model to the case of stochastic growth of the mutant, where we consider (i) a linear birth process for the mutant and (ii) a random relative growth rate for the mutant. We stress that in the following computations, the growth rate is not constant, but can have an arbitrary form. The last section is devoted to a discussion of possible extensions of this work and conclusions. An appendix, containing a straightforward mathematical derivation is included in order to make this article self sufficient.

## II. DETERMINISTIC MODEL

### A. Equal growth of WT and mutant

Consider a culture of bacteria growing from size *N*_0_ to size *N*. The growth curve can be as general as possible assuming that no death event takes place. Let *X*_*n*_ be the random variable describing the occurrence of a mutant when the WT population increases from *n* to *n* + 1. Throughout this paper, we use the term mutant to designate an individual which acquire a trait (e.g. phage or antibody resistance) which will be tested once the growth is stopped.

Denoting the mutation *probability* by *υ*,

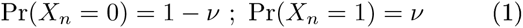

This may seem to be an approximate description, because if during a cell division, a mutation has occurred, obviously the number of wild type cells has not increased from *n* to *n* + 1. A more precise formulation would be Pr(*X*_*n*_ = *k*) = (1−*υ*)*υ*^*k*^, *i.e.,* the number of mutants when the WT population increases by one unit is geometrically distributed. However, as *υ* ≪ 1 (*υ* is usually of the order of 10^−8^), we will use the relation (1) to describe the random variable *X*_*n*_. The generalization to the geometric distribution of *X*_*n*_ is straightforward (see appendix 1). Note that most formulations of LD distribution use the above approximation.

We assume in this subsection that both mutant and the wild type population follow a deterministic, equal growth. We do not assume the growth rate to be a constant. As the mutants are similar to wild types, a mutant appearing in one copy at WT population size *n* will contribute *N/n* to the number of mutants when the wild type population reaches size *N* (figure 1). In other words, the proportion of the number of this mutant to the number of WT population will remain constant.

**Figure 1:**
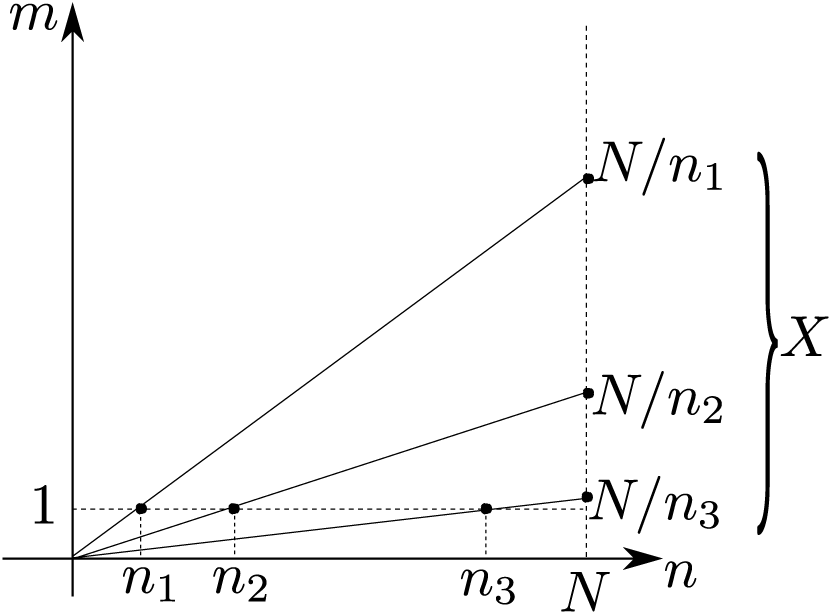
The number of mutants *X* at population size *N* is the sum of the contributions of mutants appearing at population size *n*_*i*_. The *i*–th mutant, appearing at size 1 at WT population *n*_*i*_ will be present at size *N/n*_*i*_ in the final population.

Let 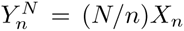 be the contribution of this mutant to the final number of mutants *X*, when the WT population reaches size *N*. Then

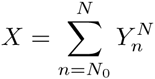

As the mutant occurrences are independent random variables, the moment (mgf) and cumulant generating functions (cgf) are

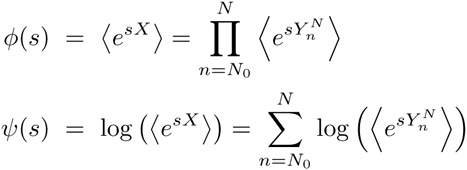

On the other hand, by its very definition,

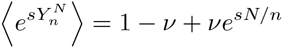

which gives the cgf as

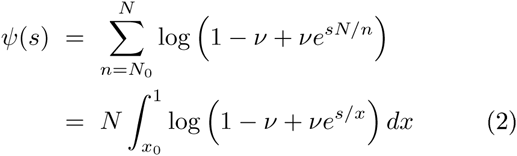

where in the second expression, we have used the continuous approximation for the sum, *x* = *n/N* and *x*_0_ = *N*_0_*/N*.

Note that this derivation is analogous to the filtered Poisson process derivation used when the problem is formulated in real time ([8],eq. 14). However, because the problem is formulated in terms of WT population size, the propagator is simply a straight line regardless of the growth rate function (figure 1).

Expanding the expression (2) to the first order in *ν* and restricting the domain of definition to *s* ≲*-x*_0_ log *υ*,

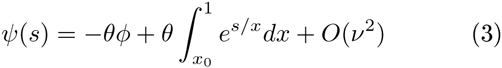

Where *θ* = *N ν* and *ϕ* = 1 − *x*_0_. For the particular case of exponential growth, the expression (3) for the cgf has been obtained by Crump and Hoel [5] and in closed form by Zheng ([8],eq.14). Note that for Zheng, *N* = *exp*(*βt*), *i.e.* the initial number of WT bacteria is 1.

The first two cumulant coefficients *κ*_*p*_ = *Ψ*^(*p*)^(0) are then

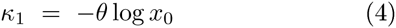

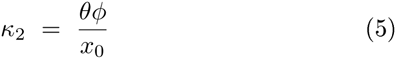

These expressions for the average (*κ*_1_) and the variance (*κ*_2_) have been obtained originally by LD for the exponential growth of bacteria [1]. As we see here, the hypothesis of exponential growth is superfluous and the expressions (4-5) are valid for arbitrary growth curves. To the leading order in *ν* and *x*_0_, the general expression for cumulant coefficients is

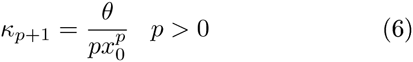

These expressions are known for the special case of exponential growth ([8],eq.9 for equal growth rate).

### B. Different growth of WT and mutant

Let us now consider the case where WT (*n*) and mutants (*m*) (once they have appeared) have similar but different growth rates:

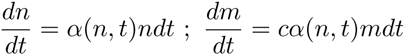

where *c* is a constant. We do not specify any particular form for the growth rate, but we suppose that the mutant follows the same law as the WT, within a constant multiplicative factor. This is the case for example where the resources are depleted by the growth of the bacteria, and the mutant is inferior to the WT for its duplication.

Time can be eliminated between the above equations: *dm/dn* = *c*(*m/n*). A mutant appearing at one copy when the population size is *n* will contribute (*N/n*)^*c*^ to the final number of mutants (figure 2). The computation of the preceding section can be repeated and leads to

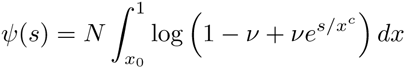

**Figure 2:**
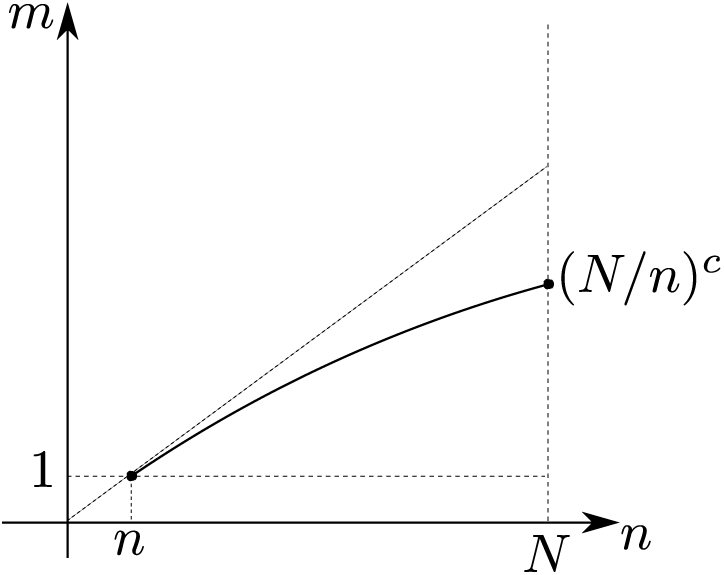
Contribution of mutants with different growth rate.

Keeping only the leading term in *υ* and *x*_0_, for *c*≠1*/p*, the *p-*th cumulant coefficient is given by

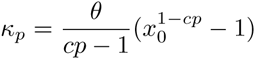

The case *c* = 1*/p* can be recovered from the above formula by taking the limiting value for *c* → 1*/p* and reads

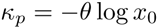

For the particular case of exponential growth (*α* = *Cte*), this is the expression given by Zheng ([8],eq.9).

### C. Variable mutation probability

In most models the mutation probability *ν* is considered to be a constant and independent of time, *i.e.,* population size. This is a sound hypothesis when the mutation involves only point-mutations on the chromosome. However, traits such as virus or antibiotic resistance may involve many point mutations before the trait is functional. Bacteria at the end of the growth process, having achieved more divisions, may be more prone to mutate to the given trait than bacteria at the early stage of the growth. A crude approximation of the above phenomena will be a mutation rate that depends on the population size *υ* = *υ*(*n*). The formulation for the number of mutants of the previous section does not suppose a constant rate of mutation and the relation 2 for *Ψ*(*s*) remains valid. The first two cumulants are given by

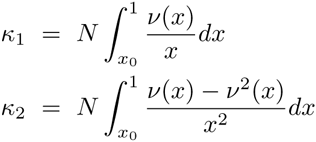

## III. STOCHASTIC GROWTH OF MUTANTS

### A. General discussion

Until now, we have considered the deterministic propagation of the mutant from its appearance at population size *n* to its final value at population size *N*. We will denote the propagator as 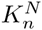. The contribution of the mutant appearing at WT population size *n* to the final population *N* was expressed as

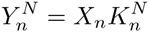

For the deterministic case, 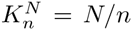 and the mgf of 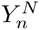 was simply

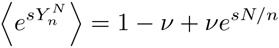

We will now consider for the mutant a stochastic propagator 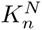 (figure 3). Because *X*_*n*_ takes only the values 0 or 1,

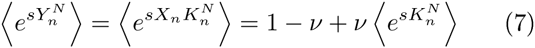

**Figure 3:**
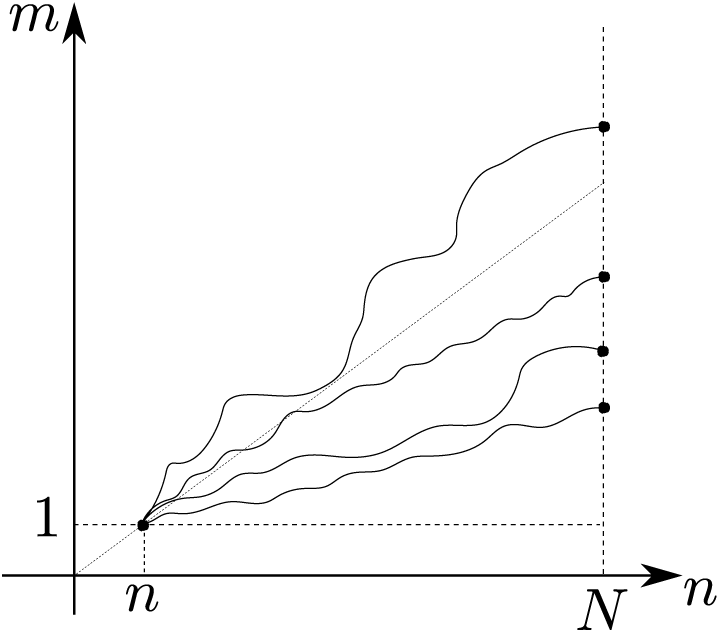
The stochastic propagator 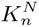 of the mutant appearing at WT population size *n*

(see appendix 2). Therefore, all the discussion of the preceding section naturally generalizes to stochastic propagation and the cgf for the number of mutants at population size *N* is

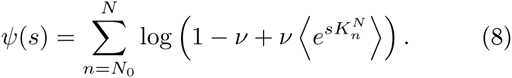

Knowing the statistical properties of the propagator gives access directly to the statistical properties of the total number of mutants. Before applying this concept to specific cases, let us compute the first two cumulant coefficients:

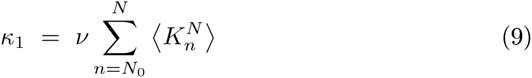

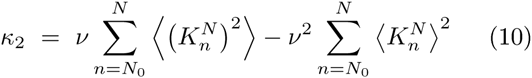

The mean *κ*_1_ is what we already had in the deterministic case, where 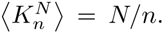 Let us express the second moment of 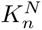 as a function of its mean and variance 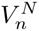

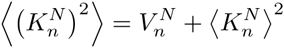

.Then

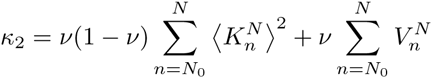

The first term on the RHS of the above relation is what we already had in the case of deterministic growth. The second term is the contribution of the stochasticity of the propagator to the variance of the number of mutants at population size *N*.

### B. Linear birth process

Consider the case where the growth of the WT is deterministic and continuous

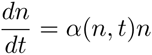

while the mutant, once it has appeared, follows a stochastic growth with transition probability density

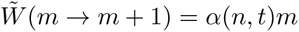

where 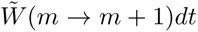 is the probability that this lineage of the mutant has increased its size by one unit in the interval [*t, t*+ *dt*[. This model was first introduced by Lea and Coulson[2] for exponential growth case *α*(*n, t*) = *α*.

A note of caution should be made here. Although widely used, the linear birth model may not be very realistic, as bacterial division times are not exponentially distributed. Indeed, after a division, a bacterium needs to elongate again to its original size before being able to divide again, so that the next division time cannot be smaller than a finite time *τ*. In fact, the distribution of division times around the time *τ* is fairly narrow and the division process is much less random than a linear birth process. The phenomenon has been experimentally investigated by a microfluidic device by Wang et al [17]; the overestimation of the mutation rate by a linear birth model, for the exponential growth case, has been investigated by Ycart [11].

Let us now come back to the linear birth model. As in the previous section, the real time is not the best choice of independent variable and we can write the stochastic growth of the mutant in terms of WT population size: noting *W* (*m → m* + 1)*dn* the probability that a mutant has divided when the WT population size ∈ [*n, n* + *dn*[, we have

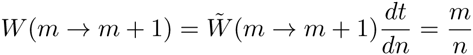

The master equation governing the growth of the mutant is

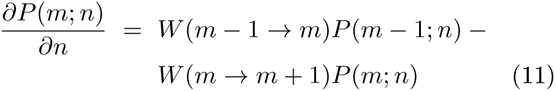

and the mgf of the propagator 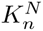 is given by (see appendix 3):

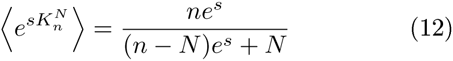

Using now the expression (8) and the continuous variable *x* = *n/N*, we obtain the cumulant generating function of the number of mutants

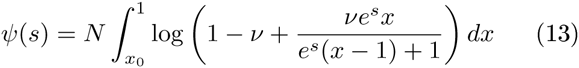

Note that the above integral can be expressed in an analytical, albeit cumbersome, form. The expressions for the two first cumulant coefficients are

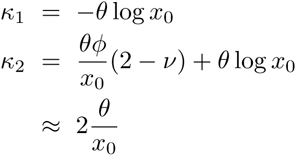

Where as before, *θ* = *N ν* and *ϕ* = 1 *− x*_*0*_. The variance of the number of the mutants is now approximately twice what we had for the deterministic case. The exponential growth case can be recovered from the above expression and is equal to the expressions given by Lea and Coulson and Zheng ([8], eq.52-53).

Other cumulant coefficients can be readily recovered by multiple derivation of expression (13). Restricting the computations to the leading order of *ν* and *x*_0_, the expression for the cumulant coefficients is (see appendix 4)

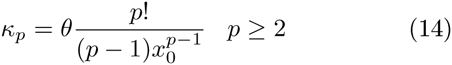

Note that even in the case of exponential growth, no general expression for the cumulant coefficients could be obtained by classical methods ([8]). Comparing the above expression with the deterministic case where *κ*_*p*_ = 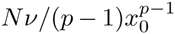, we see that indeed the linear birth process induces large amplification of the *p-*th cumulant coefficient by a factor of *p*!.

*The probabilities.* To compute the probabilities, it is more advantageous to use the probability generating function (pgf)

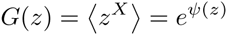

where *Ψ*(*z*) is defined from (13) by setting *z* = *e*^*s*^, *i.e.*:

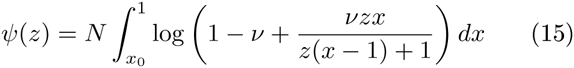

Approximating the above expression to the first order in *υ*, we have

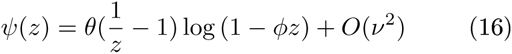

where *θ* = *N ν* and *ϕ* = 1 − *x*_0_. We therefore obtain a simple expression for the probability generating function

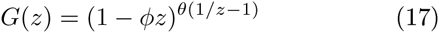

For the case of exponential growth, the above expression (without the *ϕ* factor) was first discovered by Lea and Coulson[2]; omitting the *ϕ* factor however results in divergent moments. The correct expression can be seen in the Zheng review ([8], eq 65). We stress again that relation (17) is very general and does not depend on the assumption of exponential growth. Note that in the case of exponential growth, *θ ∼* exp(*βt*) and all the moments diverge as *t* ⟶ ∞. This divergence, discussed by Bartlett and later by Zheng [8], cannot be cured within the framework of the exponential growth model. No such divergence exists in the present formulation, as the WT population size, following any realistic growth curve, will remain finite.

In order to evaluate the probabilities, we have first to compute *Ψ*^(*p*)^(0). Expanding expression (16) in powers of *z* we have

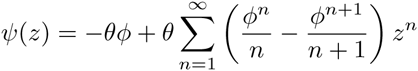

Keeping only the first leading terms in *θ* and *x*_0_ we have

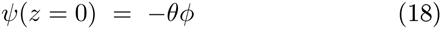

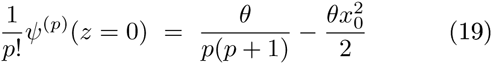

where the second term in 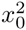 can be neglected for *p* ≪ 1*/x*_0_ and *θϕ ≈ θ*. We can now use the Faà Di Bruno Formula [18] to compute the *k–*th derivative of *G*(*z*) at *z* = 0 and obtain the probabilities (see appendix 5):

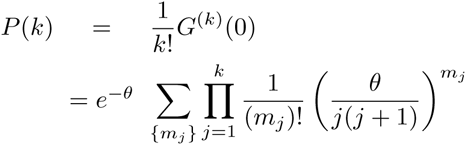

where the sum is taken over all *n-*tuples {*m*_*j*_} such that 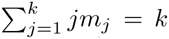. The above formula can be easily programmed to compute numerically the probabilities. The only linear term in *θ* in the above formula is for *m*_*k*_ = 1, *m*_*i*<*k*_ = 0. Therefore, for *θ* ≪ 1, the probabilities take the simple form of

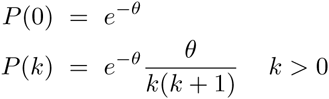

The explicit expression for the probabilities in the linear birth model has been intensely investigated by many authors, and reviewed by Zheng ([8], 5.3) and the above expressions are known for the constant linear birth model.

Experimentally, obtaining the probabilities implies a very large number of parallel cultures and the above expressions may not be of great practical use.

*Different growth rate.* The discussion of the preceding subsection can be extended to take into account a different relative growth rate for the mutant compared to the wild type:

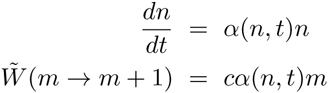

Repeating the discussion of the preceding sections, the cumulant generating function in this case is (see appendix 3)

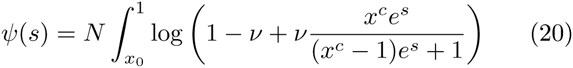

from which the cumulant coefficients can be deduced as before.

### C. random relative growth rate

The traditional formulation of LD distribution assumes that all mutants are similar in their growth function. As many different mutations can bring a bacteria to the same phage resistance, this assumption may seem too restrictive and can be easily relaxed. For example, a comprehensive study of this phenomenon has recently been published [19] where the growth rate of *all* mutants in the gene TEM-1, conferring resistance to the antibiotic cefotaxime, where measured and shown to be variable in some conditions.

Consider the case where the growth of both mutants and WT are deterministic as in subsection II B

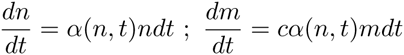

but the relative growth rate *c* is now a random variable (figure 4): when it appears, a mutant picks a relative growth rate *c* from a given distribution, which is transmitted to its progeny.

**Figure 4:**
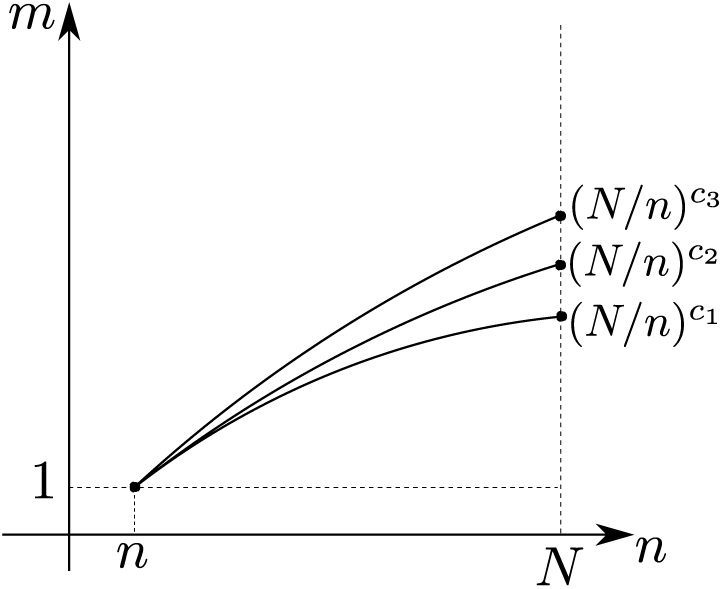
Deterministic growth, random relative growth rate.

Following the discussion of subsection III A, the propagator this time is

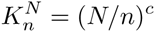

where *c* is now a random variable. Let us denote *ρ*(*s*) with its moment generating function

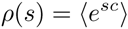

Then according to relations (9-10), the first two cumulants are now, to the first order in *υ*:

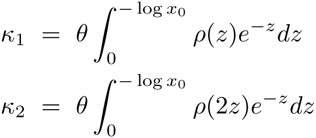

For example, if *c* follows a Normal distribution *c* = *𝒩* (*μ, σ*), then *ρ*(*s*) = exp(*μs* + (1*/*2)*σ*^2^*s*^2^) and these expressions can be evaluated in terms of the error function. For the specific case of *μ* = 1 and *σ* ≪ 1, restricting the computation to the leading orders of *ν* and *x*_0_, we have

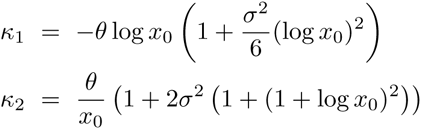

## IV. DISCUSSION AND CONCLUSION

In the previous sections we have given the general solution of the LD distribution through the derivation of its cumulant generating function. The key point to this derivation is to change the independent variable from time to WT population size, which is indeed the relevant variable whatever the growth function, a mutation can occur only when a WT cell divides and WT population size changes from *n* to *n*+1. This consideration considerably simplifies the solution of the problem and allows us to extend the solution to arbitrary growth curves for the WT population. The mathematical formulation applies equally well to the case of deterministic and stochastic growth.

This mathematical formulation is sufficiently simple to allow for many generalizations, some of which have been considered in this article: for example variable mutation probability (subsection II C) or random relative growth rate (subsection III C) have been investigated..

Other generalizations that we have not developed can be considered. For example, for the stochastic growth case, we have only considered the linear birth process. Other, more realistic cases can be envisaged where the distribution of the division times are not exponential, along the lines developed by Ycart[11]. One can also consider the case where both mutants and WT grow stochastically. These generalizations would be straightforward if the moment generating function of the propagator 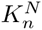 can be derived in explicit form. Another possible generalization would be to take into account the experimental uncertainty on the initial and final value of the WT population, and its influence of the estimation of mutation probability, as has been considered by Ycart and Veziris[20].

To summarize, we have developed in this article a versatile method for investigating the Luria Delbriück distribution with an arbitrary growth function. The method uses only very few measurable parameters, namely the initial and final number of the WT population. We believe that the method we propose here can be used as a simple basis for further investigations of the LD distribution.

## Appendix: Mathematical details

### 1. Geometric distribution of mutants appearing at WT population size *n*

We have noted *ν* the probability of apparition of a mutant when *a* wild type cell divides, and *X*_*n*_ the random variable tracking the number of mutants appearing when WT population changes its size from *n* to *n* + 1. The probability of producing *k* mutant during this change is geometrically distributed:

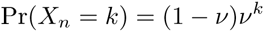

As *υ* ≪ 1, we have approximated *X*_*n*_ by a binary process (*X*_*n*_ = 0, 1) in the article. This constraint can be relaxed. Following notations of subsection II A,

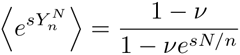

and

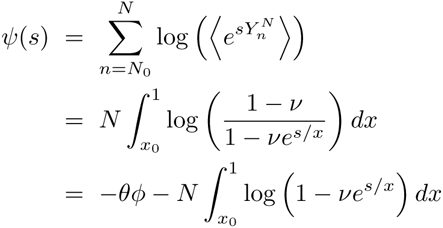

where *θ* = −*N* log(1 − *ν*) and *ϕ* = 1 − *x*_0_. The above expression is equal to expression (2) to the first order in *υ*.

### 2. Moment generating function of 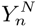 in the stochastic case

Consider the random variable 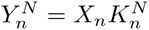 where *X*_*n*_ is a boolean variable *P*(*X*_*n*_ = 0) = 1 − *ν* and *P*(*X*_*n*_ = 1) = *ν* (subsection III A); For simplicity, we suppose that 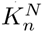 is a positive discrete random variable. Then,

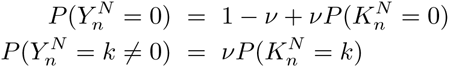

The mgf is then

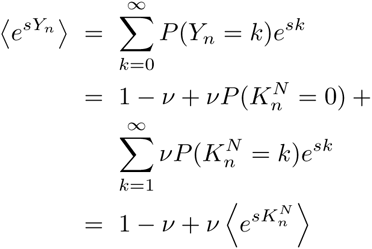

which is the expression (7).

### 3. Moment generating function of the propagator 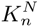

A mutant appearing in one copy when the WT population size is *n*_0_ will reach size 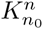 when the WT population reaches size *n*. The forward master equation governing the probabilities of the propagator, 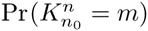 is given by equation (11). The mgf of the propagator, *ϕ*(*s, n*) therefore obeys the following equation:

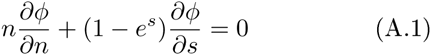

with the boundary conditions

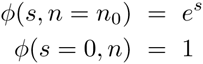

Equation (A.1) is a linear first order partial differential equation that can be solved by the method of characteristics:

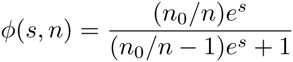

Changing now the notation to denote *n* as the WT population size of the mutant occurrence, and *N* the final size of the WT population, we obtain the expression (12).

If the relative growth rate of the mutant is not 1 but *c*, where *c* is an arbitrary constant, the transition probability for the mutant once it has appeared is

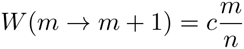

and *ϕ*(*s, n*) obeys the following equation:

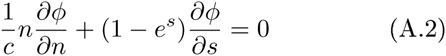

This equation can be transformed into equation (A.1) by a simple scaling *n* ⟶ *n*^*c*^ and the mgf is therefore given by

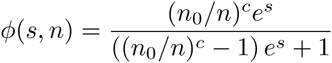

from which the cumulant generating function of the number of mutants can be deduced.

### 4. Cumulant coefficients for the stochastic growth

To the first order in *ν*, the cgf for the linear birth model (subsection III B) is given by

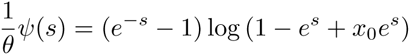

Expanding the above function into powers of (1 − *e*^*-s*^), we have

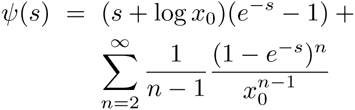

Evaluating the *p - th* derivative at *s* = 0, we have

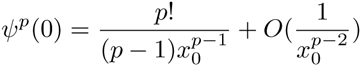

which is the expression given in equation (14).

### 5. Probabilities

The Faà Di Bruno Formula for chain derivation of the function *f*(*g*(*z*)) is[18]

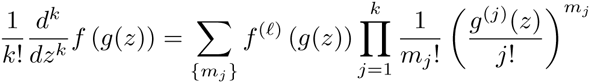

where the sum is taken over all {*m*_*j*_} such that

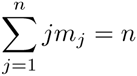

and 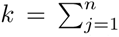. In the present case, *g*(*z*) = *Ψ*(*z*) and *f* (*u*) = exp(*u*). Specifically, *g*(0) = −*N ν* = −*θ*, and therefore *f*^(ℓ)^(*g*(0)) = exp(−*θ*). On the other hand, according to relation (19),

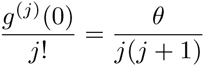

## Acknowledgements

I sincerely thank Marcel Vallade, Olivier Rivoire and Erik Geissler for the critical reading of the manuscript.

